# High-throughput Single Cell Motility Analysis using Nanowell-in-Microwells

**DOI:** 10.1101/2025.06.18.660403

**Authors:** Pan Deng, Wenze Lyu, Deasung Jang, Kerryn Matthews, Simon P. Duffy, Hongshen Ma

## Abstract

Cell motility is important to many biological processes including cancer, immune response, and tissue repair. Conventional assays measure bulk cell motility, potentially overlooking important heterogeneity and missing important high motility subpopulations. Here, we introduce a high-throughput single-cell motility assay using nanowell-in-microwell plates to precisely track single cell position and analyze their migratory trajectories. By physically confining individual cells in nanowells, we eliminate cell-cell interactions and simplify cell segmentation and tracking. Using this platform, we characterized the motility of single cells across different culture conditions to identify distinct motility phenotypes. Single-cell trajectory analysis revealed pronounced directional persistence, with cells predominantly maintaining their direction of travel and trajectory along nanowell boundaries. Additionally, our approach facilitates the generation of labeled image datasets suitable for AI models to rapidly identify cell motility phenotypes from single-cell images. Together, our platform provides a robust, scalable method to analyze cell motility phenotypes and migration behavior at single-cell resolution.

**Significance Statement:** A novel single-cell motility assay that uses confinement in nanowell-in-microwell to enable high-throughput profiling of single cell motility phenotypes.

## Introduction

Cell migration is a critical part of many biological processes including development, angiogenesis, wound healing, and cellular immunity. Cell migration is also a key part of cancer progression, associated with invasion and metastasis. Therefore, cell migration assays are valuable tools for identifying therapeutic agents and evaluating drug efficacy^1–3^. Traditional cell migration assays count the number of cells that transit from an occupied space to an empty space, which may be formed between two chambers in a transwell assay, or in a cell-free gap on agar in a wound healing assay^4–6^. These assays provide a convenient measurement of the mean migration rate across a cell population. However, primary human tissues, including tumor digests, constitute mixed cell populations with heterogeneous phenotypes. Even cells from the same cell line can exhibit different behaviors due to variations in signaling and protein expression^7,8^. For these samples, the bulk migration rate may obscure the behavior of rare but clinically important cell subpopulations. For example, Zhou and colleagues reported on compounds that appear to inhibit tumor cell migration by reducing the migration of most cells, thus lowering the average migration rate^9^. Yet, these compounds were ineffective against a minority subpopulation of fast-moving cells that retain invasive and metastatic potential. This finding underscores the critical need for granular assays capable of analyzing cell motility at the single-cell level.

Significant progress has been made to develop microfluidics technologies to analyze single cell migration by measuring the distance cells travel in confined microchannels^9–15^. These microfluidics assays have demonstrated predictive power for cancer patient outcomes^13^ and show promise as tools for identifying novel clinical biomarkers^16^. Despite their potential, these methods require precise loading of individual cells into separate microchannels and alignment to a common starting point^17^. Both of these steps are technically challenging to implement and difficult to scale for high-throughput experiments. Furthermore, microchannel-based assays restrict cell migration analysis to a single dimension, oversimplifying cell motility and potentially missing biologically meaningful patterns of movement. An alternative strategy is to use continuous microscopy to track the migration of single cells on imaging substrates^18,19^. Although this analysis can be multiplexed by rapidly cycling the microscope stage across multiple fields of view, this approach is limited by errors introduced by the presence of proximal cells or cell aggregates, which can obscure single cell trajectories and confound analysis^20,21^.

Here, we present a novel strategy that simplifies the challenge of cell tracking and enables high-throughput profiling of single cell migration. This strategy involves monitoring movement of individual cells confined within nanowells, where the spatial isolation of each cell eliminates interference from neighboring cells. To ensure compatibility with existing high-throughput imaging platforms, we fabricate a high-density nanowell array inside standard 384-well microwell imaging plates. Cells are deposited into each microwell and randomly distributed into nanowells based on the Poisson distribution. Time-lapse microscopy is then used to capture the position of individual cells within each nanowell over time. Confinement in nanowells not only simplifies cell segmentation but also yields a large dataset of high-quality single-cell images suitable for training deep learning models.

## Results

### Experimental Approach

To simplify single cell tracking and enable high-throughput single-cell motility assay, we fabricated nanowell-in-microwell plates using a previously described process^22^. Briefly, open-top nanowells measuring 70×70×60 µm (*l*×*w*×*h*) arranged in rectangular arrays were fabricated using photolithography on a glass slide substrate (**Fig. 1A**). The patterned glass slides were then bonded to a standard 384-well plastic well plate frame, resulting in ∼1,200 nanowells in each microwell. Cells were seeded into each microwell at a density corresponding to ∼30% of the total number of nanowells in each microwell to maximize single-cell occupancy, based on Poisson statistics (**Fig. 1B**). After seeding, cells were cultured for two days to promote adhesion and acclimatize to the glass substrate. For motility analysis, cells were fluorescently labeled using Calcein Green (**Fig. 1C-D**) and then imaged using automated microscopy to track cell position in each nanowell every hour for 12 hours (**Fig. 1E**).

**Fig. 1.**
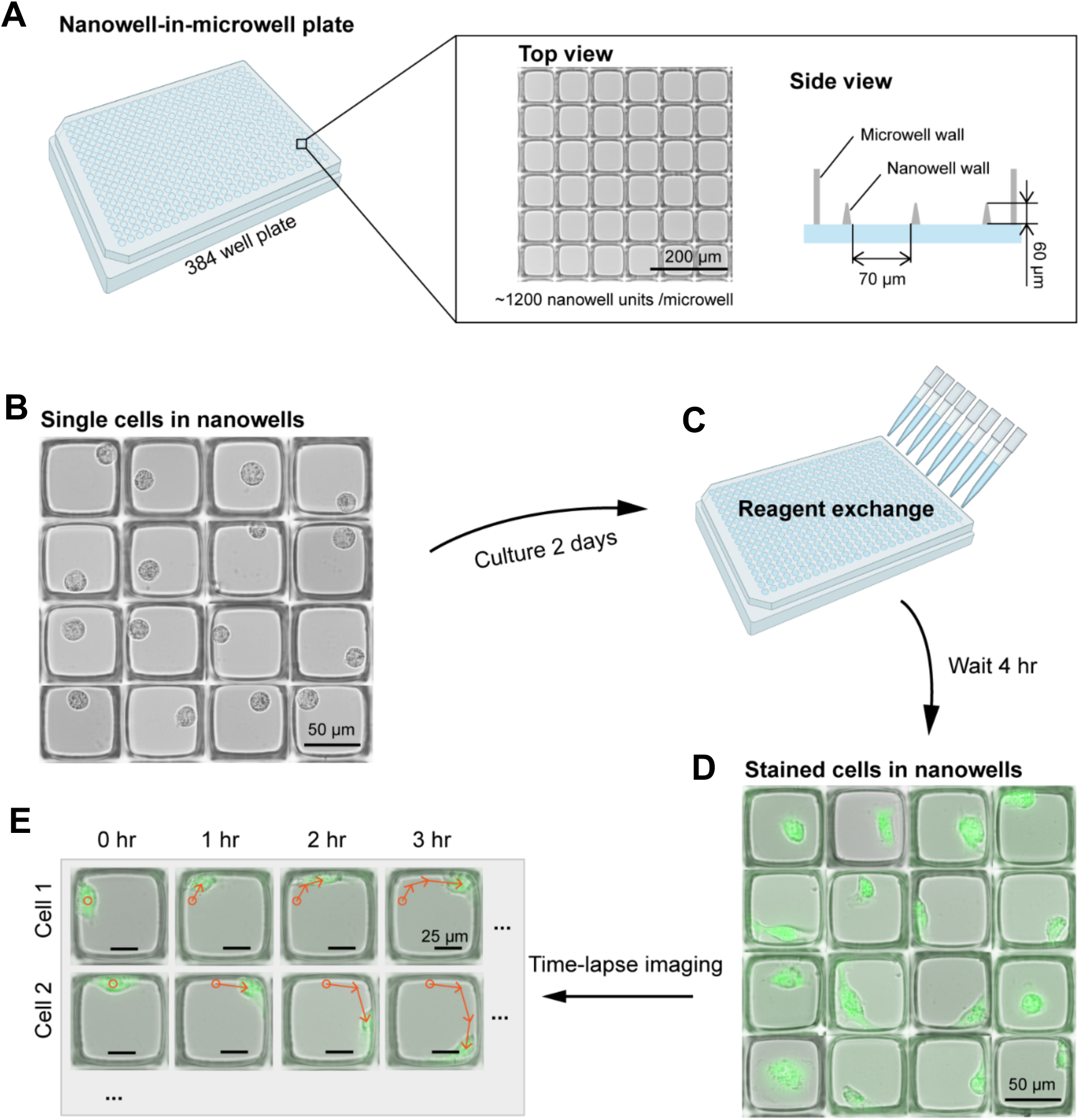
Single cell motility analysis using the nanowell-in-microwell platform. (**A**) Nanowell-in-microwell plate: A 384-well plate that contains ∼1,200 nanowells, where the dimensions of each nanowell are 70×70×60 µm (*l*×*w*×*h*). (**B**) To perform single cell motility analysis, cells are randomly seeded into nanowells and cultured for two days to enable cell adhesion. (**C**) The culture medium is refreshed with different treatments. (**D**) The cells are stained in the nanowells with Calcein AM and imaged at 1-hour intervals for 12 hours. (**E**) The images are analyzed to track cell trajectories.

### Image Analysis Pipeline

To analyze single-cell motility from time-lapse microscopy images (**Fig. 2A**), we developed an imaging processing pipeline to analyze brightfield and fluorescence images to track the position of single cells within individual nanowells. This pipeline consisted of a series of steps including: (a) nanowell segmentation, (b) live cell detection, (c) nanowell image filtering, (d) single cell position and morphology analysis in individual nanowells. Nanowell segmentation was performed by using user interface software to overlay a 2D grid on the brightfield images of nanowells to determine the range of coordinates for individual nanowells (**Fig. 2B**). Typically, 1,024 single nanowell images (242×242 pixels) can be segmented from each microscopy field. Each segmented nanowell is assigned a unique address based on pixel coordinates to analyze across experimental time points. After segmentation, we performed a series of filtering steps to eliminate invalid nanowells. First, we analyzed each nanowell to identify live cells based on the Calcein AM fluorescence signal to eliminate nanowells that lacked live cells (**Fig. 2C**). Next, we estimated the number of viable cells in each nanowell to eliminate nanowells that contained more than one cell (**Fig. 2D**). Finally, we analyzed the full time-series images for each nanowell to eliminate nanowells that showed a loss of cell viability or a change of cell count (*e.g.* resulting from cell division) over the 12-hour analysis window (**Fig. 2D-F**). For each remaining nanowell image, we analyzed the fluorescence image to extract the centroid position of each cell, as well as the length of each cell measured across its longest axis (**Fig. 2G-H**). The change in centroid position is used to determine the average motility of each cell, while the change in cell length is used to determine the average elongation rate of each cell.

**Fig. 2.**
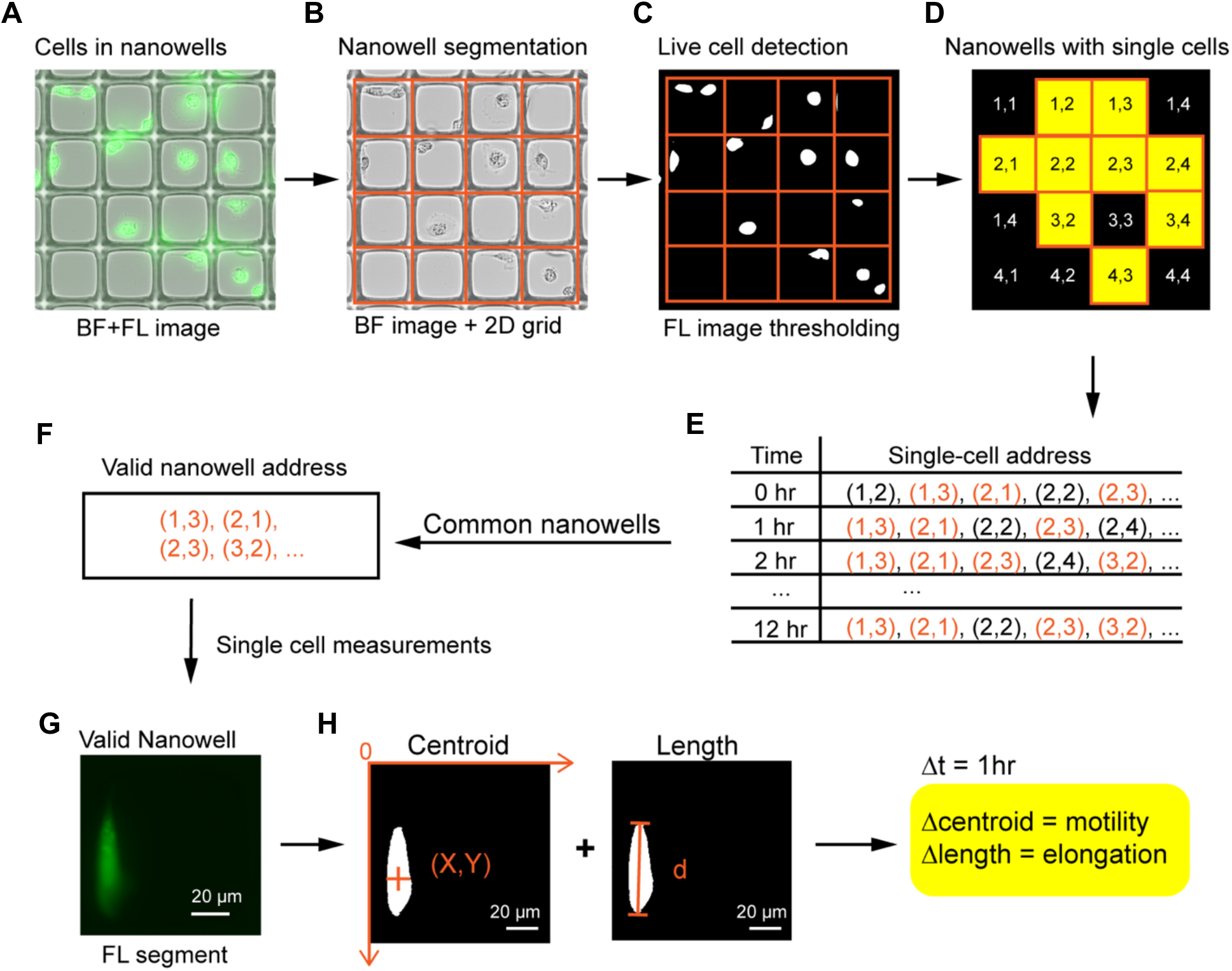
Data analysis pipeline. (**A**) Acquire brightfield (*BF*) and fluorescence (*FL*) images of cells in nanowells. (**B**) Segment individual nanowell images by mapping a 2D grid onto the brightfield images. (**C**) Identify live cells by thresholding fluorescent images. (**D**) Identify nanowells occupied by single cells. (**E-F**) Analyze the nanowells at multiple time points. Only nanowells that consistently detected with a single cell are deemed valid. (**G-H**) The cell motility and elongation rate in valid nanowells are tracked by thresholding the fluorescence image to assess the cell centroid and length.

### Single Cell Motility Analysis

MDA-MB-231 breast cancer cells are known to exhibit genetic and phenotypic heterogeneity^21^. We characterize the migration behavior of MDA-MB-231 cells at the single cell level under three conditions designed to induce distinct motility phenotypes. Specifically, serum starvation (0% FBS) produced low-motility cells^23^, standard culture (10% FBS) produced cells with normal motility, and TNF-α stimulation (10% FBS + 10 ng/mL TNF-⍺) produced high-motility cells^24,25^. We tracked the centroid positions of 300 single cells for each condition at one-hour intervals over 12 hours. We found that serum starvation (0% FBS) resulted in a mean motility of 4.4±3.1 µm/hr, standard culture cells resulted in a mean motility at 12.2±9.6 µm/hr, while TNF-⍺ stimulation resulted in a mean motility at 15.9±11.0 µm/hr (**Fig. 3A**).

**Fig. 3.**
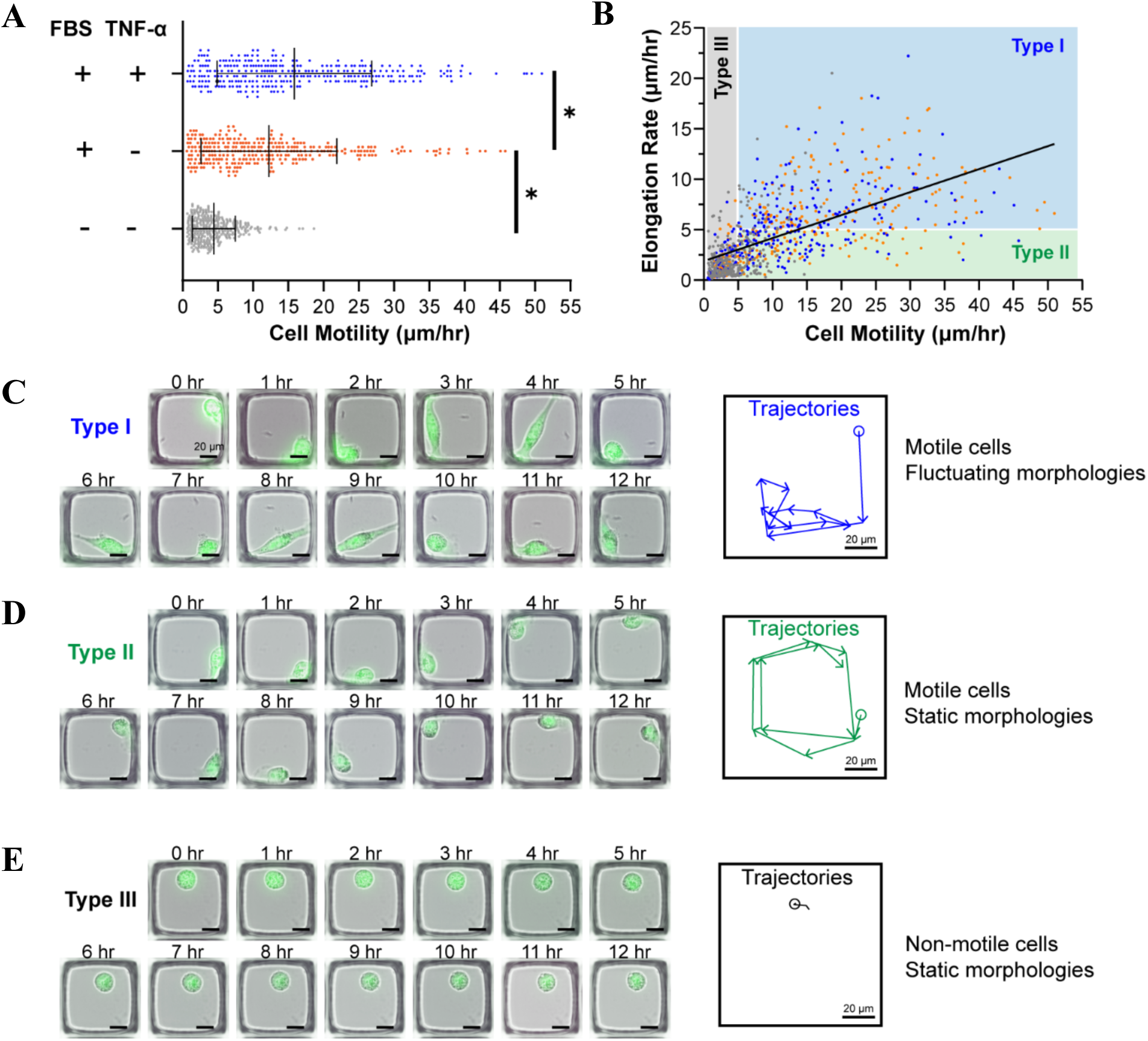
Cell motility and morphology analysis. **(A)** Cell motility under starvation culture (0% FBS), normal culture (10% FBS), and stimulated culture (10% FBS, 10 ng/ml TNF-α). The data for each condition is collected from 300 single cells in 6 microwells. *p<0.001 **(B)** Cell motility versus elongation rate for all three conditions (grey: starvation culture; orange: normal culture; blue: stimulated culture) categorized into three types. The solid line is the linear fit of the scatter plot (Pearson’s r=0.6, P<0.0001). **(C-E)** Time-lapse images of representative single cells acquired hourly showing differences in morphology (left) and trajectories (right).

Single-cell profiling of motility revealed considerable heterogeneity in the motility phenotype among individual MDA-MB-231 cells. This heterogeneity was reflected by the wide deviation around the motility for each culture condition (**Fig. 3A**). To further characterize heterogeneity in cell motility, we defined a motility threshold of 5 µm/hr and assessed the proportion of cells from each culture condition that exceeded this threshold. In serum-starved cells, only 33% of cells exceeded this threshold motility rate, while most of the cells propagated in standard culture conditions (75%) and TNF-α stimulation culture (83%) exceeded this threshold motility rate. These findings suggest that serum-rich and TNF-α stimulation conditions not only enhance the migration rate of already motile cells but also increase the overall proportion of motile cells in the population.

A defining feature of migrating cells is the formation of protrusions that elongate the cell along the direction of motion^26^. To investigate whether cell elongation correlates with motility, we define the elongation rate as the average magnitude of the cell length difference between consecutive one-hour intervals over a 12-hour period. Analyzing cell elongation rate versus average migration speed for all 900 single cells over the three conditions, we found that elongation rate was correlated with migration speed in each condition, with an overall Pearson’s r=0.6 (**Fig. 3B**). Based on this analysis, we can categorize cell movement into three distinct phenotypes: Type I cells with characteristic motility and elongation rate exceeding 5 µm/hr, Type II cells showing motility with lower cell elongation rate, and Type III cells representing non-motile cells (**Fig. 3C-E**). This analysis underscores how single-cell nanowell profiling can associate specific cellular morphologies with motility phenotypes to reveal functional subpopulations.

### Single Cell Trajectory Analysis

Beyond measuring motility, our image-based single-cell profiling enables tracking of single cell migration paths. We analyzed the trajectories of Type І and Type II motile cells with motility >20 µm/hr to study cell behavior during migration. Specifically, we measured the turning angle (θ) at each time point (*t*), which is defined as the angle between the cell trajectory from *t-1* to *t hr* and the cell trajectory from *t* to *t+1 hr* time points (**Fig. 4A**). An acute angle (|θ| < 90⁰) indicates forward movement, while an obtuse angle (|θ| > 90⁰) indicates a reversal in the direction of movement. Across all time points, 93% of movements were forward (|θ| < 90°), suggesting cells generally persisted in their current direction.

**Fig. 4.**
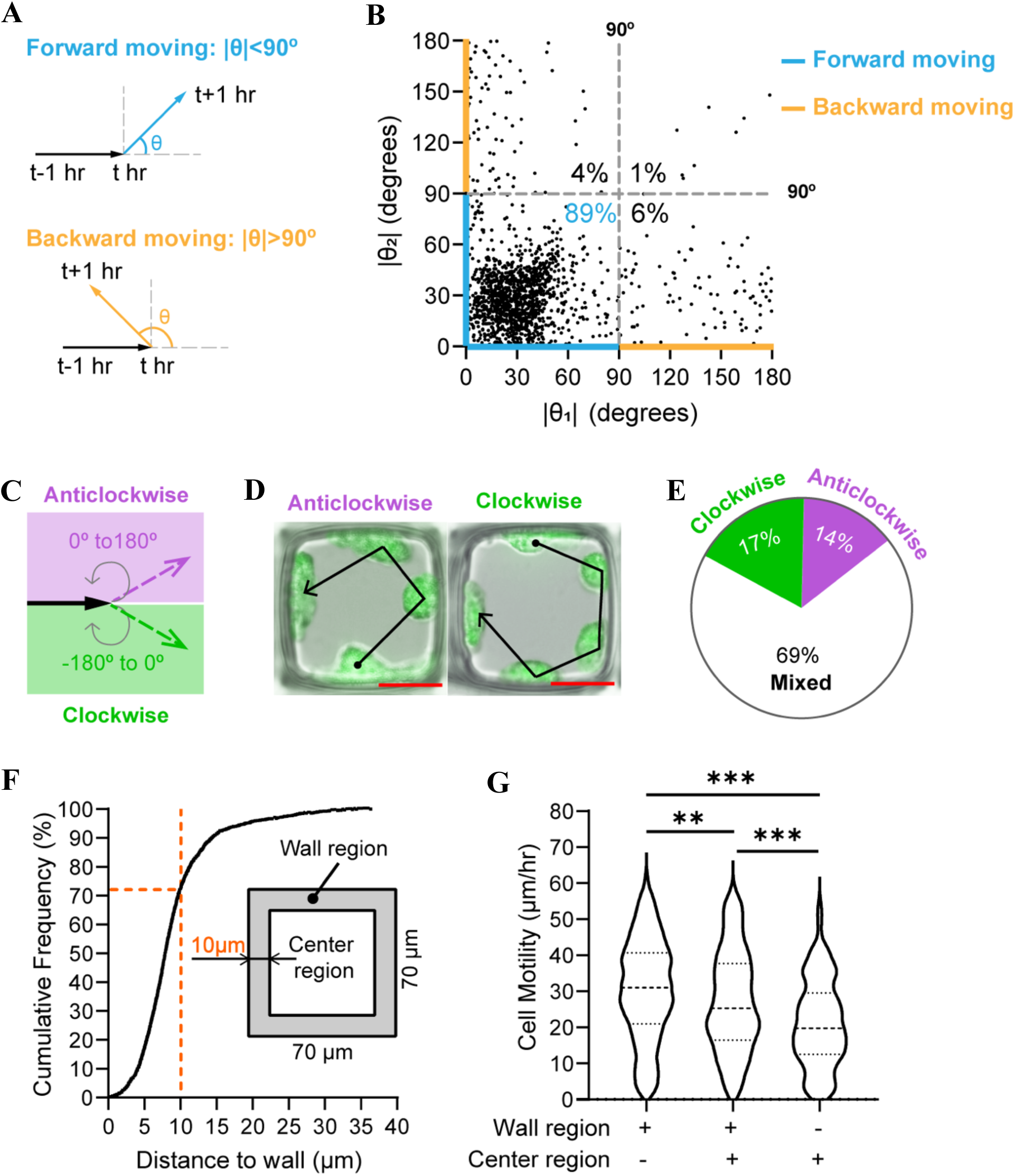
Trajectory analysis of highly motile cells (motility > 20 µm/hr). **(A)** The turning angle (|θ|) at *t* hr is determined by the movement from *t-1* hr to *t+1* hr. Cell trajectories are considered to be forward moving if |θ| < 90°; and backward moving if |θ| > 90°. **(B)** Relationship between two consecutive turning angles (|θ₁| and |θ₂|) measured after each time point (N=1490). **(C)** Definition of clockwise (−180°<θ< 0°) and anticlockwise (0°<θ<180°) cell motion based on turning angle θ. **(D)** Representative images of single cells exhibiting exclusively clockwise or anticlockwise turning angles. Scale bar = 20 µm. **(E)** Distributions of three different migrations over the 12 hours. Anticlockwise: cells exhibited only anticlockwise turning angle; clockwise pattern: cells exhibited only clockwise turning angle; mixed pattern: cells exhibited both anticlockwise and clockwise turning angles. **(F)** Cumulative frequency of the distance between cell centroids to the nearest edge of the nanowell at every time point. Inset: definition of center and wall regions, which are almost equal in area. **(G)** Motility when cells are confined to the nanowell wall region only, the center region only, and transitioning between the two regions (N ≥ 130/group). All data were analyzed using a *t*-test. **p<0.01, ***p<0.001

To assess whether past movements influenced subsequent migration patterns, we analyzed consecutive turning angles from *t* to *t+1 hr* time points, which we define as |θ₁| and |θ₂| (**Fig. 4B**). We found that cells moving in a forward direction continued along that forward trajectory in 89% of cases. After a cell reversed its trajectory from forward movement to backward movement, the cell continued along the new direction at the next time point in 88% of cases. These results indicate that cells primarily persist in their most recent direction and are minimally influenced by prior trajectory.

To analyze how the Type І and Type II motile cells interact with nanowell walls, we tracked cell trajectories over a 12-hour period. At each time point, we consider a turning angle (θ) between 0° and 180° as the cell trajectory turning in the anticlockwise direction, and a turning angle between −180° and 0° as a cell trajectory turning in the clockwise direction (**Fig. 4C-D**). To characterize the overall rotational behaviour of each cell, we analyzed the full sequence of turning angles across the entire 12-hour period. This analysis revealed that 31% of cells exhibited consistent rotational motion, with 17% showing exclusively clockwise turning and 14% showing exclusively anticlockwise turning (**Fig. 4E**).

Finally, we analyzed the spatial distributions of high motility cells in their nanowell over 12 hours. Each nanowell was divided into two regions of equal area: a wall region (10 µm band near the nanowell edge) and a center region (**Fig. 4F**). We observed a strong bias for cell migration along the wall region, where cells were detected within this region in 73% of images. Additionally, cells migrating exclusively along the wall exhibited the highest motility, while those confined to the center showed the lowest motility (**Fig. 4G**). These findings suggest that when encountering a barrier, cells preferentially move along it to maintain their directional persistence.

### Using Deep Learning to Predict Cell Motility

Our nanowell-in-microwell platform enables imaging and precise tracking of thousands of single cells in nanowells isolated from other cells. We can leverage this capability to train a deep learning model to identify cell motility phenotypes without performing time-laps microscopy experiments. We performed this study by selecting nanowell images captured at the time point where the cells that moved >5 µm were considered motile, and cells that moved <5 µm were considered idle. To train this classification model, we implemented a convolutional neural network (CNN) with four convolutional layers (3 × 3 kernel size), three max-pooling layers, two dense layers (50 neurons each), and a final fully connected output layer (**Fig. 5A**). We collected a dataset of 15,594 labeled images for training the CNN, distinguishing between actively migrating and idle cells (**Fig. 5B**). Model performance was assessed using six-fold cross-validation, yielding an average validation accuracy of 82%, demonstrating consistency across different subsets of the dataset. When tested on previously unseen cells, the model achieved 80% accuracy in predicting the actively migrating states and 82% accuracy for idle states (**Fig. 5C**), indicating that deep learning could effectively infer cell motility states from static images. To interpret the features driving the model’s predictions, we generated saliency maps (**Fig. 5D**), which revealed that the network primarily focused on cell boundaries. This suggests that morphological differences, particularly cell shape, play a key role in distinguishing motile cells from idle cells.

**Fig. 5.**
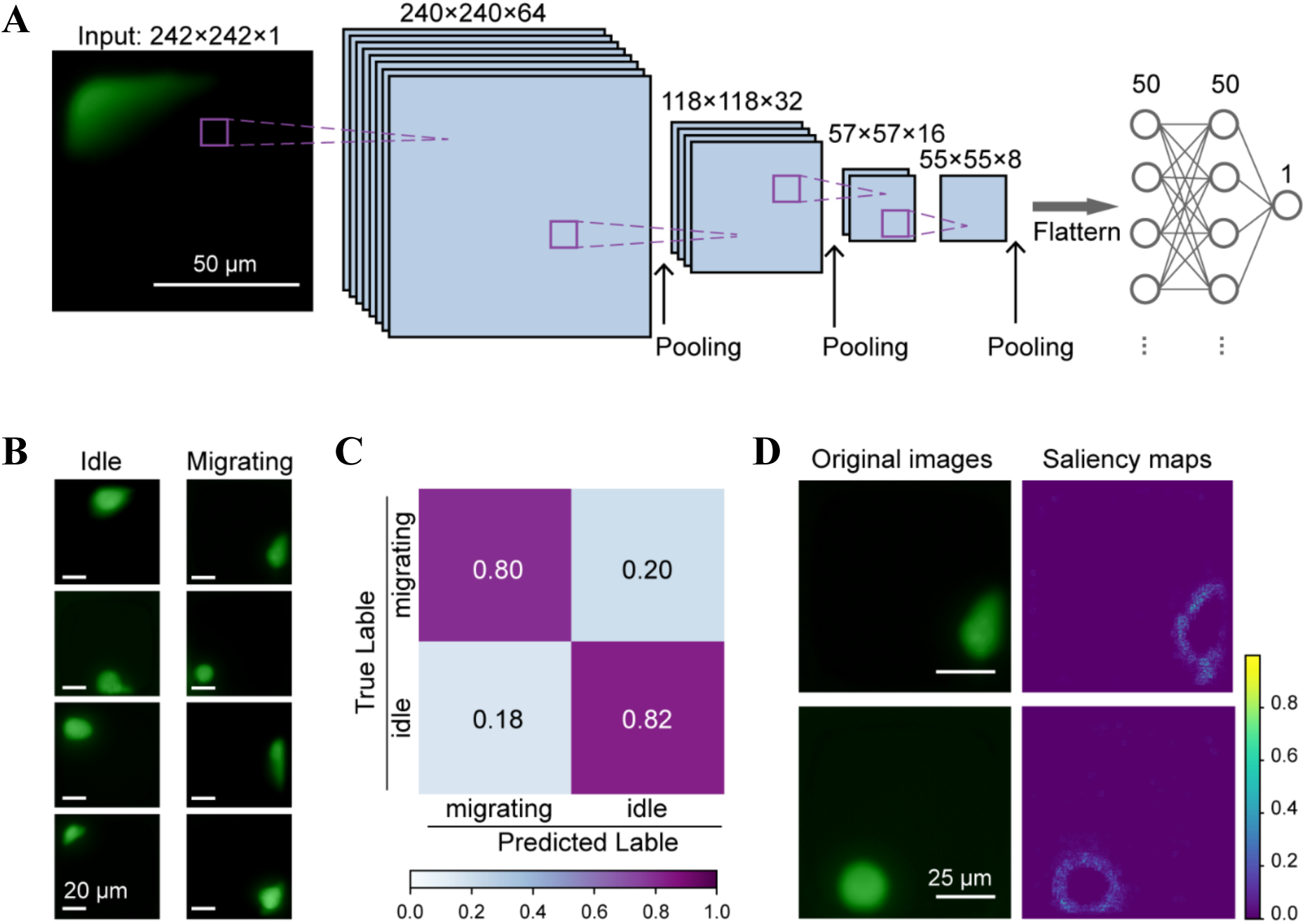
Predicting cell motility using deep learning. **(A)** Architecture of a convolutional neural network model used to predict cell motility. **(B)** Example fluorescence image of cells in the inert state and motile state. **(C)** The confusion matrix obtained from evaluating our model on previously unseen test cell images. **(D)** Saliency maps of two randomly selected cells.

## Discussion

In this study, we leveraged nanowell-in-microwell plates to develop a high-throughput single-cell motility assay to enable detailed characterization of motility phenotypes within heterogeneous cell populations. Traditional assays measure cell motility at the bulk level, potentially obscuring rare subpopulations with distinct migratory behaviors^4–6,9^. While approaches using microfluidic^9–15^ and continuous microscopy^18,19^ can offer single cell resolution, these methods are limited by technical challenges associated with introducing cells into microchannels, aligning cells to a common starting point, or separating cells migrating in crossing paths, all of which severely limit throughput. Our approach addresses this issue by confining single cells in nanowells, which allow them to be isolated from each other and characterized using time-lapse microscopy.

The key innovation of our approach is to confine single cells in nanowells to eliminate the bottleneck associated with cell tracking and interference from adjacent cells. Integrating nanowell arrays in ANSI standard 384-well imaging plates, which provides compatibility with existing high-throughput microscopy and automated liquid-handling workflow. Each microwell contains ∼1,200 individual nanowells, which enables simultaneous tracking of hundreds of isolated single cells.

We demonstrated the effectiveness of our platform by identifying motility phenotypes of MDA-MB-231 cells under various culture conditions, revealing significant heterogeneity within each population. Specifically, cells cultured under serum starvation exhibited low motility, while standard culture conditions and TNF-⍺ simulation produced higher proportions of migratory cells. Even though confinement within nanowells limited the physical space available for migration, our measurements of cell migration speed were similar in range to those observed by cell tracking by continuous microscopy in standard microwell substrates^27–29^.

By leveraging single cell imaging, our assay also provided new insights into cell migration behavior. We observed strong directional persistence with 93% of cells maintaining continuous forward migration. Furthermore, when cells bumped up against the side walls and changed their trajectory, significant fractions of cells maintained a consistent turning direction (clockwise or counterclockwise) over the 12-hour observation period. These observations align with known cytoskeletal mechanisms governing migration that involve a commitment to actin polymerization at the leading edge of the cell^30,31^. Our assay can distinguish true changes in cell motility rate from shifts in net migration caused by variations in forward-backward movement frequency or trajectory changes, which is a capability that is not achievable using microchannel-based microfluidic assays. We also noted that cells predominantly migrated along the nanowell periphery (73% of tracked time points), with the highest motility observed in cells confined to the wall region. This may be associated with their strategy to maintain the direction persistence and mechanosensing that is known to enhance cell migration on curved substrates^32,33^. By comparing cell migration throughout the nanowell, it may be possible to elucidate conditions that alter cell sensitivity to this mechanosensing phenomenon.

Beyond its immediate application in cell behavior analysis, our nanowell-based approach generates a rich dataset that can be leveraged for deep-learning analysis. Historically, deep learning in microscopy has been primarily applied to challenges such as cell segmentation and tracking, which are major bottlenecks for real-time analysis in live-cell imaging^34^. By simplifying cell tracking through nanowell confinement, we achieved a higher throughput, allowing us to generate sufficient data to train a deep learning model capable of distinguishing migrating versus idle cells. In contrast to existing models that require multi-frame time-lapse data^35–37^, our approach provides a streamlined method for inferring cell motility from static snapshots. Future work will focus on expanding the training dataset to improve model generalizability and robustness across diverse cell types and treatment conditions. The integration of deep learning with high-throughput motility assays has the potential to provide an automated approach to distinguish motility phenotype directly from a single microscopy image.

In summary, our nanowell-in-microwell single-cell motility assay overcomes the cell-tracking bottlenecks of conventional continuous microscopy methods. By confining cells within nanowells in a standard 384-well plate format, we ensure compatibility with automated liquid handling and imaging systems. This platform enables both velocity and trajectory measurements at high throughput, producing datasets suitable for deep-learning analysis of cell motility phenotypes. The potential applications of this platform extend beyond basic research in phenotypic characterization to clinical drug testing, where it could be used to identify and target rare, highly motile tumor cell subpopulations during drug screening. Furthermore, our motility dataset could support the development of future AI-driven models to predict cell migration and therapeutic efficacy directly from cell morphology without requiring motility assays. Together, these advancements position our nanowell-based approach as a tool for studying cancer cell motility and developing targeted therapies.

## Methods

### Cell Lines and Reagents

MDA-MB-231 cells (ATCC number: HTB-26) were cultured in Dulbecco’s modified Eagle medium (DMEM, Gibco) with 10% (v/v) heat-inactivated fetal bovine serum (FBS, Gibco) and 1% (v/v) penicillin at 37°C in 5% CO_2_. After the cells reached 75% confluence in T75 flasks (Thermo Scientific), the cells were harvested and cultured inside nanowell-in-microwell plates (200 cells/microwell) for two days to attach to the nanowell surface. Prior to motility assay, cells were conditioned for 4h in one of three media conditions: (1) DMEM with 0% FBS, (2) DMEM and 10% FBS and (3) DMEM with 10% FBS and 10 ng/mL of TNF-⍺.

### Nanowell-in-Microwell Device Fabrication

Nanowells-in-microwell devices are fabricated using a previously described process^22^. Briefly, glass slide substrates (75×50×0.3 mm, Abrisa Technologies) was rinsed first with acetone (Sigma-Aldrich) and then with ethanol (Sigma-Aldrich), before plasma cleaning. The glass slides were then treated with a solution of 10% v/v TMSPMA (M6514, Sigma-Aldrich) in ethanol for 2 hours at 70 °C followed by washing with ethanol and baking at 80 °C for one hour. The nanowell microstructures were fabricated by UV photolithography using a chrome photomask. After photolithography, the uncured polymers are removed by rinsing with isopropyl alcohol, resulting in a bas-relief structure of nanowells on the glass surface. The glass was then glued to the bottom of a 384-well plate frame (Grace Bio Labs) to form a nanowell-in-microwell platform. Each standard microwell contained approximately 1,200 nanowells, with individual nanowell dimensions of 70×70×60 µm (length × width × height).

### Cell Imaging and Motility Assay

MDA-MB-231 cells under various culture conditions were stained with Calcein AM using the manufacturer’s protocol (Thermo Fisher Scientific). Stained cells were subsequently seeded into the nanowells-in-microwell plate at low density and cultured for 48 hours at 37°C to allow for cell adhesion. Cells were imaged under brightfield and fluorescence imaging using a Nikon Ti-2E inverted microscope every hour for 12 hours.

### Image Processing and Analysis

A software program developed on Python 3.7 was used to align nanowell units to a 32×32 grid and perform segmentation of each nanowell image. Images of nanowells that were not occupied by single cells were excluded from analysis. To analyze the nanowell images, the fluorescent images were converted to a binary image based on a threshold determined from the background intensity level. Next, cells were labelled and counted using skimage.measure algorithms. The nanowell exhibiting a cell count of less than or more than one cell over 12 hours of imaging were excluded. The remaining cells were analyzed with Python using the scikit-image library for automated determination of cell centroid coordinates and measurement of the major axis of the cell. The centroid coordinate was used to determine cell position, while the measured major axis was to quantify cell length.

Cell movement was quantified by calculating the distance between cell centroids at consecutive 1-hour time point. The average cell motility was calculated from the mean distance moved between each time point over 12 hours. Similarly, cell elongation was evaluated by determining the difference in cell length at each 1-hour interval, with the elongation rate calculated based on the mean elongation per time point over 12 hours. The trajectory of each cell over time was determined by mapping the absolute coordinates of cell centroids to the nanowells at each time point.

### Deep Learning for Classification of Cell Motility

We generated a convolutional neural network model that was modified from the AlexNet model architecture in Python 3.7 using the Keras library in TensorFlow. Specifically, our network accepts a one-channel image input of 242×242 pixels. To extract image features, we utilized four convolutional layers with a kernel size of 3×3 each layer followed by a ReLU activation. The first, second, and fourth convolutional layers were each followed by a max-pooling layer with a size of 2. Then, the output was flattened into a one-dimensional array to achieve full connection with the latter two dense layers. Each dense layer had 50 units followed by a ReLU activation and 20% dropout. The final layer had one output node activated by a sigmoid function for binary classification. In model training, binary cross-entropy was defined as our loss function to measure the difference between predicted outcomes and actual labels. Each iteration of the network was trained on 25 epochs, with Adam optimization and a learning rate of 0.0001. After filtering out images of cells that were out of focus and cell doublets, there were 15,594 images/class to ensure even classes and non-biasing model training. The training was performed on a single computer with 64.0 GB of RAM and an NVIDIA graphics card (GeForce RTX 3090).

## Author contributions

H.M. supervised the study. H.M. and P.D. conceived the idea. P.D., W.L., and D.J. performed the experimental work. P.D. analyzed the data. All authors wrote the manuscript.

## Data Availability

Data and code for this article are available at https://github.com/Pan-De/single-cell-motility-analysis

## Acknowledgments and Funding

This work was supported by grants from the Natural Sciences and Engineering Research Council of Canada (2020-05412, 2020-00530, 590749-24). P.D. acknowledges funding from the China Scholarship Council and the Tai Hung Fai Charitable Foundation.

